# Scalable and interpretable secretion system annotation with Sismis

**DOI:** 10.1101/2025.09.09.675188

**Authors:** Martin Larralde, Florian Albrecht, Josefin Blom, Johan Henriksson, Laura M. Carroll

## Abstract

Secretion systems play critical roles in bacterial growth, survival, and pathogenesis. Genes encoding secretion system components often co-occur together as gene clusters in bacterial (meta)genomes. However, existing tools for secretion system annotation are unable to utilize genomic context to make predictions and are unable to detect secretion systems of novel architecture. Here, we present Sismis (<u>s</u>ecretion <u>s</u>yste<u>m</u> d<u>is</u>covery tool; https://github.com/lmc297/Sismis), a scalable, interpretable, machine learning-based tool, which detects and classifies single-locus secretion systems in bacterial (meta)genomes with high accuracy (test set area-under-the-curve [AUC] values of 0.71 and 0.92 for precision-recall [PR] and receiver operating characteristic [ROC] curves, respectively). When applied to ≈700k prokaryotic (meta)genomes, Sismis identifies 747, 439 total secretion systems comprising 15, 612 major secretion system families, >80% of which contain no previously known/annotated secretion systems. To facilitate further exploration of these data, we present the Sismis Atlas (https://sismis.microbe.dev/), an interactive secretion system database, which we use to identify a largely uncharacterized cluster of secretion systems with tight adherence (Tad) pili-like characteristics. Altogether, Sismis and its companion atlas enable accurate and interpretable secretion system annotation, exploration, and discovery at an unprecedented scale.

**GRAPHICAL ABSTRACT:** 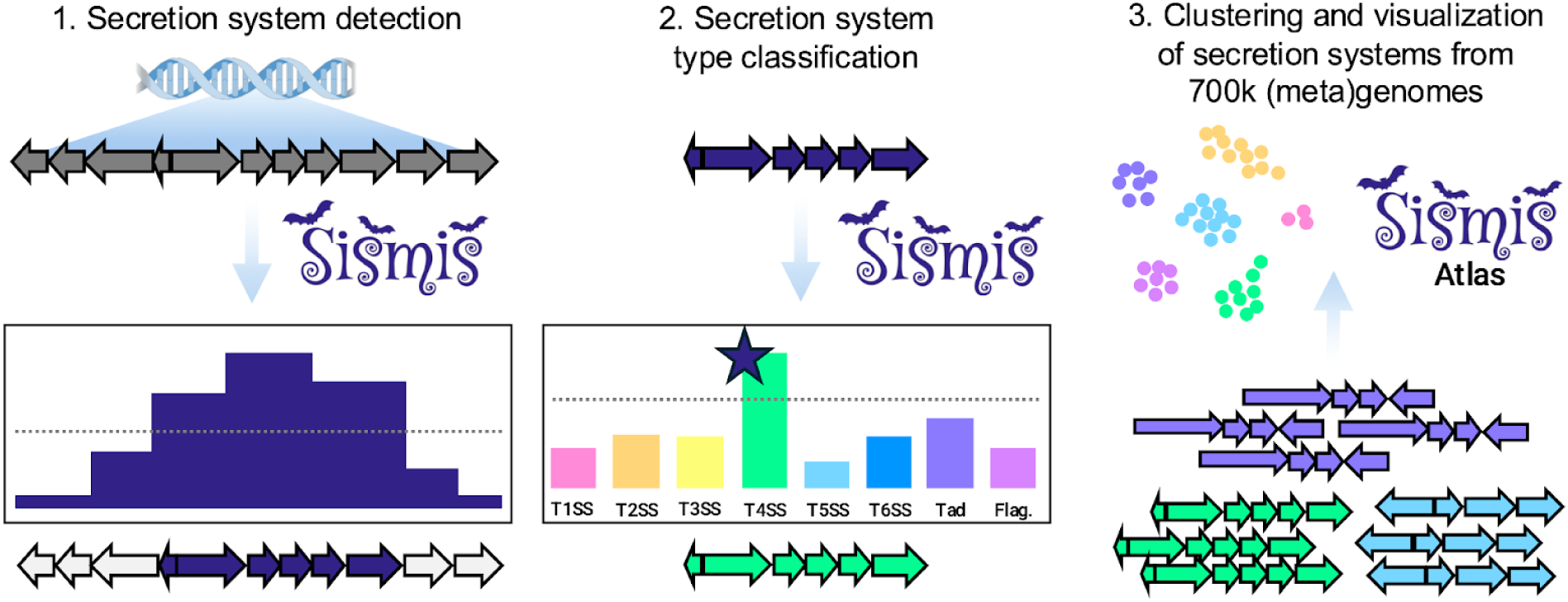

## INTRODUCTION

Bacteria utilize a range of conduits for substrate secretion, including large, membrane-spanning macromolecular complexes known as secretion systems (1–4). Secretion systems play integral roles in a wide variety of prokaryotic processes; for example, they allow bacteria to grow, survive, and adapt to changing environmental conditions, and they facilitate interactions with hosts and other organisms (including other bacteria) (1–4). Among bacterial pathogens, secretion systems play crucial virulence-promoting roles (e.g., by exporting protein toxins and effectors that directly target host cells; by promoting attachment/adhesion; via nutrient acquisition; by enabling biofilm formation) (1–5) and thus are key to understanding bacterial pathogenesis.

While their classification is not straightforward and often debated, secretion systems have been grouped into at least 11 major “types” (i.e., type I to type XI, abbreviated T1SS to T11SS, respectively) based on their composition and mechanism (3).

Historically, the discovery of novel secretion system types and subtypes has taken place in the laboratory, using experimental methods (3). However, this discovery process is often time-consuming, laborious, expensive, and limited in scope (e.g., to a selection of physically accessible strains). As such, (meta)genomic mining approaches have recently been posited as a promising alternative (3). By using computational methods to screen prokaryotic (meta)genomes for the genomic determinants of secretion systems (which often–but not exclusively–co-occur together in a single locus/as a gene cluster) (1, 3, 6), one can generate potential leads in a relatively fast and inexpensive manner, without limiting oneself to a small subset of strains (3). Thus, computational methods that can scale to the massive amounts of prokaryotic (meta)genomic data in public databases have tremendous potential to enable secretion system discovery and characterization (3).

Currently, state-of-the-art computational tools for secretion system gene cluster detection in prokaryotic (meta)genomes (e.g., MacSyFinder) are “rule-based”; they rely on manually curated sets of profile hidden Markov models (pHMMs) (7) to detect genes with known secretion system associations, and then use hard-coded “rules” to determine if a secretion system is present or not (6, 8, 9). While adequate for applications where high precision is warranted, rule-based methods inherently cannot detect secretion systems with novel architectures, and as such, likely miss novel types and subtypes that do not follow existing rules. Alternatively, sequence modeling approaches, which have been used successfully for other gene cluster detection tasks (10–13), have the potential to overcome these limitations, but thus far have not been used in the secretion system annotation space.

Here, we present Sismis (<u>s</u>ecretion <u>s</u>yste<u>m</u> d<u>is</u>covery tool; https://github.com/lmc297/Sismis), a scalable, interpretable, machine learning (ML)-based tool for *de novo* secretion system annotation. Using a combined (i) conditional random field (CRF) and (ii) random forest (RF) approach for secretion system detection and type classification, respectively, we show that Sismis detects and classifies secretion systems in bacterial (meta)genomes with high accuracy and speed.

Further, because CRFs are interpretable models, which leverage genomic context to make predictions, we look “under-the-hood” of our optimal model and identify features that the Sismis CRF deems important for secretion system detection. Finally, we use Sismis to mine the entirety of the mOTUs v3 database (i.e., nearly 700k prokaryotic [meta]genomes) (14) for putative secretion systems, and we make our predictions available to the public via an interactive atlas (i.e., the Sismis Atlas, https://sismis.microbe.dev/). Together, the methods and tools developed here facilitate large-scale exploration of the prokaryotic secretion system space.

## MATERIALS AND METHODS

### Training and test set construction

Metadata (i.e., nucleotide accession numbers, secretion system start/end coordinates, and secretion system type classifications) for all secretion systems classified as “single_locus” secretion systems were downloaded from TXSSdb (*n* = 10, 784 TXSSdb single-locus secretion system entries; http://macsydb.web.pasteur.fr/txssdb/_design/txssdb/index.html, accessed 27 September 2024) (6). Using R v4.3.2 (https://www.R-project.org/) and the ‘entrez_fetch’ command from rentrez v1.2.3 (15), all valid single-locus secretion system-containing contigs were downloaded from NCBI’s Nucleotide database (16) using their NCBI Nucleotide accession numbers (*n* = 1, 958 contigs containing 9, 756 single-locus secretion systems). The ‘stratified’ command from the splitstackshape R package (v1.4.8; https://CRAN.R-project.org/package=splitstackshape) was used to partition contigs into training and test sets, stratifying by secretion system type; this resulted in (i) a training set consisting of 1, 558 contigs with 6, 920 single-locus secretion systems (**Supplementary Table S1**), and (ii) a test set consisting of 400 contigs with 2, 836 single-locus secretion systems (**Supplementary Table S2**).

### Feature construction

The conditional random field (CRF) and random forest (RF) implemented in GECCO v0.9.10 (10) were re-trained for secretion system detection and type classification, respectively. Briefly, GECCO’s ‘gecco annotatè subcommand was used to annotate all 1, 558 training set contigs (see section “Training and test set construction” above), using the following parameters: Pfam v36.0 as the pHMMs for annotation (‘--hmm Pfam-A.hmm’) (17); a *P*-value threshold of 1E-9 (‘-p 1e-9’); unknown region masking enabled (to prevent genes from stretching across unknown nucleotides; ‘--mask’); 1 CPU (‘-j 1’). Gene and feature tables produced for each training set contig were concatenated to create a gene and feature table for the whole training set, respectively (**Supplementary Tables S3-S4**). Metadata from TXSSdb was used to create a cluster table for the training set (see section “Training and test set construction” above; **Supplementary Table S1**), and together, the training set gene, feature, and cluster tables were used for training and cross-validation (CV) of all models (described below).

### Cross validation and model selection

Hyperparameter optimization was carried out via 10-fold CV, within folds (to prevent overfitting). Briefly, training set contigs (see section “Training and test set construction” above) were shuffled using “random.shuffle” from Python v3.11.5 and divided into ten folds. For each 10-fold CV iteration, GECCO’s ‘gecco train’ subcommand was used to re-train GECCO’s underlying CRF and RF, using the following parameters: the training fold’s corresponding (i) gene, (ii) feature, and (iii) cluster tables supplied to ‘--genes’, ‘--features’, and ‘--clusters’, respectively; (iv) 28 CPUs (‘-j 28’); (v) a *P*-value threshold of 1E-9 (‘-p 1e-9’); (vi) *c2*, the L2 regularization hyperparameter, set to 0 (‘--c2 0’); (vii) *c1*, the L1 regularization hyperparameter, set to one of [0.15, 1, 10]; (viii) a sliding window size *W* (in number of genes) of one of [5, 10, 20]; (ix) a Fisher’s Exact test (FET) feature selection threshold *T* (corresponding to the proportion of features selected) of one of [0.25, 0.5, 0.75, 1]; (x) feature type set to “protein” (‘--feature-type protein’). Each resulting model was then used to detect and classify secretion systems within the respective test fold contigs, using GECCO’s ‘gecco run’ subcommand and the following parameters: (i) each test fold contig’s FASTA file supplied as input (‘-g’); (ii) the corresponding model supplied as input (‘--model’); (iii) 8 CPUs (‘-j 8’); (iv) ‘--force-tsv’ enabled (to output results for all contigs, regardless of whether secretion systems were detected or not); (v) unknown region masking enabled (‘--mask’); (vi) Pfam v36.0 as the pHMMs for annotation (‘--hmm Pfam-A.hmm’); (vii) a *P*-value threshold of 1E-9 (‘-p 1e-9’); (viii) ‘--merge-gbk’ enabled (to merge predicted secretion systems into a single GenBank; **Supplementary Dataset S1**). Based on area-under-the-precision-recall-curve (AUPR) scores (discussed below), models with *c1* = 10 and *W* = 20 were consistently among the top performers (**Supplementary Figure S1**); as such, additional values of *T* were tested, i.e., *T* = [0.05, 0.1, 0.15, 0.2], and all models were evaluated via 10-fold CV as described above (**Supplementary Figure S2**, **Supplementary Dataset S2**). All models with *c1* = 10 and *W* = 20 were additionally evaluated as described above using leave-one-type-out (LOTO CV), with CV folds corresponding to secretion system types (per TXSSdb; **Supplementary Figure S3**, **Supplementary Dataset S3**).

All models were evaluated using per-gene receiver operating characteristic (ROC) and precision-recall (PR) curves. Briefly, GECCO “.genes.tsv” files produced for each contig were concatenated into a single tab-separated (TSV) file. The GenomicRanges R package (v1.54.1) (18) was used to identify TXSSdb secretion systems that overlapped with genes in the concatenated genes TSV. Genes were assigned a value of “1” (part of a secretion system) or “0” (not part of a secretion system). The resulting file was used to construct ROC and PR curves, using the ‘roc.curvè and ‘pr.curvè functions from the PRROC package v1.3.1 (19, 20), with secretion system-(1) and non-secretion-system-(0) associated genes supplied to ‘scores.class0’ and ‘scores.class1’, respectively, and per-gene secretion system probabilities reported by GECCO (i.e., the “average_p” column; **Supplementary Figures S1-S3, Supplementary Datasets S1-S3**). AUPR scores obtained via 10-fold and LOTO CV were used to identify a final set of hyperparameters (i.e., *c1* = 10, *W* = 20, *T* = 0.15) and secretion system probability threshold (0.80), which were implemented as defaults in Sismis.

### Training and evaluation of the final Sismis model

After hyperparameter selection (see section “Cross validation and model selection” above), the final Sismis model was trained on all 1, 558 training set contigs using GECCO’s ‘gecco-vv train’ subcommand and the following parameters: the full training set (i) gene, (ii) feature, and (iii) cluster tables supplied to ‘--genes’, ‘--features’, and ‘--clusters’, respectively (see section “Feature construction” above); (iv) 28 CPUs (‘-j 28’); (v) a *P*-value threshold of 1E-9 (‘-p 1e-9’); (vi) *c2* = 0 (‘--c2 0’); (vii) *c1* = 10 (‘--c1 10’); (viii) *W* = 20 (‘-W 20’); (ix) *T* = 0.15 (‘--select 0.15’); (x) feature type set to “protein” (‘--feature-type protein’; **Supplementary Tables S1, S3-S4**). The resulting final model was then used to detect and classify secretion systems within the full 400-contig test set (see section “Training and test set construction” above), using GECCO’s ‘gecco run’ subcommand, each test set contig’s FASTA file supplied as input (‘-g’), the final model supplied to ‘--model’, and the following parameters: ‘-j 1--hmm Pfam-A.hmm--force-tsv--mask -p 1e-9 --merge-gbk’. ROC and PR curves were constructed as described above (see section “Cross validation and model selection”; **Supplementary Dataset S4**).

In addition to the full 400-contig test set, the final Sismis model was evaluated on a conservative 180-contig subset of the test set, which contained contigs with low average nucleotide identity (ANI) relative to contigs in the training set (to prevent highly similar contigs from appearing in both the training and test sets; due to e.g., duplicate database entries and/or highly similar genomes in TXSSdb). To construct this subset of contigs, FastANI v1.33 (21) was used to calculate ANI values between all contigs in the training and test sets (default settings); only test set contigs that shared < 90 ANI with all training set contigs were maintained (*n* = 180 contigs in the conservative subset; **Supplementary Dataset S5**).

### Introspection of the final Sismis model

Plots of protein domains included in the final Sismis CRF and their associated weights (i.e., protein domains/weights in the ‘model.state.tsv’ file, with “label” == 1; see section “Training and evaluation of the final Sismis model” above) were constructed using R v4.4.0 and ggplot2 v3.5.1 (22) (**Supplementary Tables S5-S6**). A Gene Ontology (GO) term (23) enrichment analysis was conducted using R and topGO v2.56.0 (24), with the goal of identifying GO terms enriched among secretion system–associated protein domains included in the final Sismis CRF (i.e., positively weighted protein domains selected using feature selection threshold *T* = 0.15, compared to protein domains from a similar CRF with no feature selection employed/*T* = 1.0). Briefly, protein domains included in the final Sismis CRF (i.e., using *T* = 0.15) with CRF weight > 0 were assigned a value of “1” (**Supplementary Table S5**), while all other protein domains were assigned a value of “0”. Protein domains included in a CRF with no feature selection threshold applied (*T* = 1.0; all other parameters identical to those in the Sismis CRF), which were not included in the Sismis CRF (*T* = 0.15) were additionally each assigned a value of “0” (representing the protein domain “universe” used for GO enrichment; **Supplementary Table S7**). Pfam protein domain-to-GO term mappings were downloaded (https://current.geneontology.org/ontology/external2go/pfam2go, version date 2024/04/08 20:52:48, accessed 12 February 2025) and reformatted for compatibility with topGO (**Supplementary Table S8**). The ‘readMappings’ command from topGO was used to load protein domain-to-GO mappings into R; GO terms without protein domains and protein domains without GO terms were removed. For each GO ontology (i.e., biological process [BP], molecular function [MF], and cellular component [CC]), a new topGO object was created:

‘‘‘

bp <-new(“topGOdata”, # create new object ontology=[one of “BP”, “MF”, or “CC”] # specify one of BP, MF, or CC allGenes=crf.binary, # binarized vector of secretion system-associated domains annotationFun=annFUN.file, file = “terms.tsv”, # domain-to-GO mappings nodeSize=3) # set the node size to 3

’’’

Each topGO object was supplied to topGO’s ‘runTest’ function and used to perform a Fisher’s exact test (‘statistic=”fisher”’), using the ‘weight01’ algorithm (to account for GO graph topology; ‘algorithm = “weight01”’). Protein domains with *P* < 0.05 were considered to be significant, with no additional corrections applied (‘weight01’ accounts for multiple testing; https://bioconductor.org/packages/release/bioc/vignettes/topGO/inst/doc/topGO.pdf, accessed 5 June 2025, **Supplementary Table S9**).

### Speed and memory usage benchmarking

To assess the extent to which feature selection improved computational performance, we applied (i) the final Sismis model (with a reduced feature space, i.e., feature selection threshold *T* = 0.15; see section “Training and evaluation of the final Sismis model” above) and (ii) a CRF with identical hyperparameters except *T* = 1.0 (i.e., no feature selection) to a diverse set of prokaryotic genomes and compared per-genome speed and memory usage. Briefly, each CRF was used to mine all species representative genomes from the proGenomes v3 database (25) (*n* = 41, 777 genomes), using GECCO’s ‘gecco run’ subcommand, each genome as input (‘-g’), 1 CPU (‘-j 1’), and the following parameters (see section “Cross validation and model selection” above for a detailed description): ‘--mask --hmm Pfam-A.hmm -p 1e-9 --merge-gbk’. Nextflow v24.04.2.5914 (26) was used to log compute time and memory usage via the ‘tracè option (**Supplementary Table S10**). Plots were constructed using R v4.4.0, ggplot2 v3.5.1, reshape2 v1.4.4 (27), and gridExtra v2.3 (https://CRAN.R-project.org/package=gridExtra).

### Construction of the Sismis Atlas

The mOTUs v3 database (14) was downloaded using the “motus_genomes_download” script (https://github.com/motu-tool/motus_v3_genomes/blob/main/motus_genomes_download, accessed 12 October 2023; *n* = 694, 542 [meta]genomes). Each (meta)genome was then queried for secretion systems using the final Sismis model (see section “Training and evaluation of the final Sismis model” above; **Supplementary Dataset S6**). The resulting secretion system gene clusters from Sismis (*n* = 747, 439 .gbk files) were concatenated alongside single-locus secretion systems from TXSSdb (*n* = 9, 756 GenBank files downloaded from NCBI as described previously) (28). The resulting concatenated GenBank file (*n* = 757, 195 total secretion system gene clusters) was supplied as input to IGUA v0.1.0 (28), which was used to delineate gene cluster families (GCFs) using average linkage hierarchical clustering, a clustering distance of 0.80, and 72 CPUs (**Supplementary Dataset S7**).

A UMAP was constructed to visualize the resulting GCF representative secretion systems (*n* = 15, 612; **Supplementary Dataset S7**). Briefly, to create a two-dimensional visualisation of the dataset, the GCF representative secretion systems were annotated based on their Pfam protein domain content (v35.0), as described previously (28), to generate a protein domain composition matrix. The visualization itself, a UMAP, was subsequently created in R (v4.3.1) using the packages Seurat (v5.0.2) (29), Signac (v1.12.0) (30), and reticulate (v1.42.0; https://CRAN.R-project.org/package=reticulate). To load the protein domain composition matrix into R, the Python library scipy was accessed through a custom Conda environment (Python v3.11.4, Scipy v1.15.1) (31).

Thereafter, the function ‘load_npz()’ was used to import the composition matrix. The composition matrix was converted to a chromatin assay using the function ‘CreateChromatinAssay()’, followed by a conversion to a SeuratObject (‘CreateSeuratObject()’, with the assay options set to “peaks”). The matrix underwent normalization using the ‘RunTFIDF()’ function prior to feature selection (‘FindTopFeatures()’, with the ‘min.cutoff’ set to “q0”). Subsequently, singular value decomposition was used for dimensional reduction (‘RunSVD()’), and the top 30 dimensions were used as input for two-dimensional UMAP dimensional reduction (‘RunUMAP()’, with the reduction set to “lsi”). The function ‘FindNeighbors()’ was then used to compute a nearest-neighbor graph, using 30 UMAP dimensions and the reduction set to “lsi”. To select an appropriate resolution for the clustering, a clustree was generated for resolutions between 0.10 and 0.90 with 0.05 intervals, using the package clustree (v0.5.1; **Supplementary Figure S4**) (32). Thereafter, the clusters were generated with the ‘FindClusters()’ function, with the resolution set to 0.30 and the smart local moving algorithm selection (algorithm = 3; **Supplementary Table S11**). To cluster the UMAP at a higher resolution (for tight adherence [Tad] subcluster investigation; discussed in detail below), the ‘FindClusters()’ function was re-run with the same settings, except the resolution was set to 0.50 (**Supplementary Table S12**). Note that, for reproducibility, the function ‘set.seed()’ was used during UMAP construction.

### Investigation of tight adherence (Tad) subclusters

Investigation of two tight adherence (Tad) subclusters within the Sismis Atlas UMAP was carried out using R v4.4.0 and the following packages: Seurat v5.2.1, topGO v2.56.0, ggplot2 v3.5.1, gridExtra v2.3. Briefly, the ‘FindMarkers’ function from Seurat was used to identify protein domain markers of two Seurat clusters within the Sismis Atlas UMAP (i.e., Seurat clusters 4 and 31 in the UMAP constructed at resolution = 0.5; see section “Construction of the Sismis Atlas” above), using the Wilcoxon Rank Sum test (‘test.use = “wilcox”’, the remaining settings were set to their defaults; **Supplementary Table S13**). Seurat’s ‘FeaturePlot’ function was used to overlay individual protein domains on the Sismis Atlas UMAP (default settings; **Supplementary Figure S5**).

A topGO GO enrichment analysis was conducted using protein domains and associated raw *P*-values produced by Seurat’s ‘FindMarkers’ function as described above (see section “Introspection of the final Sismis model”), but with the following changes: (i) for each ontology (i.e., BP, MF, CC), a topGO object was created using a vector of raw *P*-values produced by Seurat’s ‘FindMarkers’ function, ordered from smallest to largest (supplied to the ‘allGenes’ argument), with “selected” protein domains corresponding to those with Seurat adjusted *P*-value < 0.05 (supplied to the ‘geneSel’ argument); (ii) each topGO object was supplied to topGO’s ‘runTest’ function and used to perform a Kolmogorov-Smirnov test (‘statistic=”ks”’), using the ‘weight01’ algorithm (to account for GO graph topology; ‘algorithm = “weight01”’) and increasing score order (‘scoreOrder = “increasing”’; **Supplementary Table S14**).

Individual secretion systems in Seurat cluster 31 were investigated by extracting secretion systems and 50 Kbp flanking regions from their source contig using BioPython v1.84 (33). To ensure quality, only Seurat cluster 31 secretion systems from complete (i.e., closed) genomes located within 50 Kbp of contig edges were considered. The resulting Seurat cluster 31 secretion system regions were annotated with Prokka v1.14.6 (34) as follows (where ‘cluster’ corresponds to the secretion system ID produced by Sismis and ‘region’ corresponds to the secretion system region extracted from the source contig): ‘prokka --outdir “${cluster}_prokka_results” --kingdom Bacteria --cpus 1 --addgenes --locustag locus --centre X --compliant --prefix “${cluster}” “${region}”’. The resulting GenBank files were compared and visualized alongside secretion systems from TXSSdb using clinker (v0.0.12, default settings; **Supplementary Figure S6**) (35).

## RESULTS

### Sismis detects secretion system gene clusters with high accuracy and speed

Sismis employs a two-stage approach for secretion system detection and type prediction, respectively, inspired by approaches used previously for biosynthetic gene cluster detection and classification (**Figure 1A**) (10, 12). Briefly, in the first stage, secretion system-encoding gene clusters are detected in a user-supplied prokaryotic (meta)genome, using ordered vectors of Pfam protein domains as features and a conditional random field (CRF), which classifies each gene as secretion system-encoding or non-secretion-system-encoding (**Figure 1A**). In the second stage, secretion system gene clusters identified by the CRF (if any are present) are passed to a random forest (RF) classifier; using each secretion system’s Pfam protein domain composition as features, the secretion system is then classified into one or more of eight single-locus secretion system types defined by TXSSdb (T1SS-T6SS, Tad, and/or Flagellum) (6) or “Unknown” (if a secretion system type cannot be assigned with confidence; **Figure 1A**).

**Figure 1.**
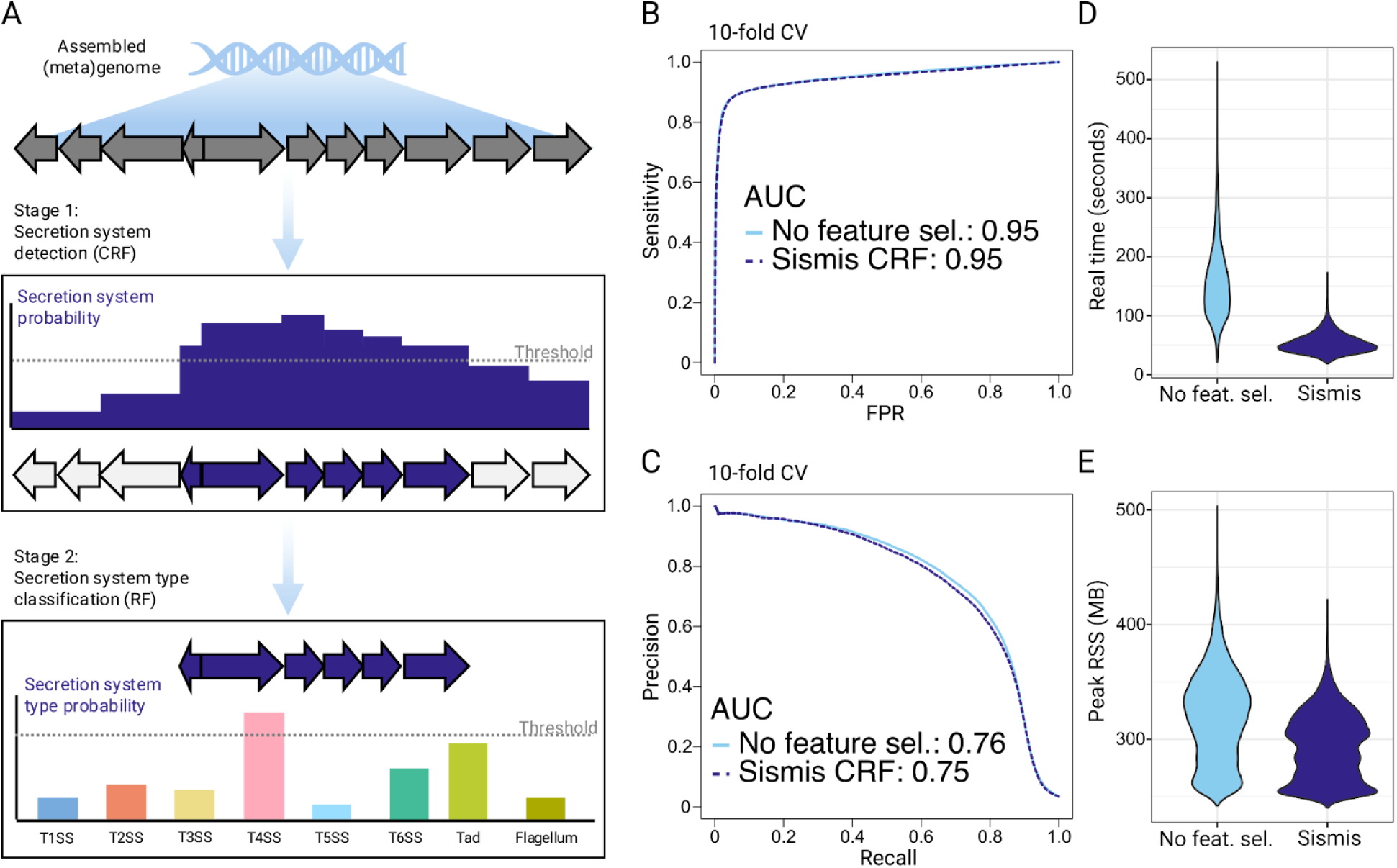
A reduced feature space enables fast, accurate secretion system detection. **(A)** Overview of the Sismis workflow. In the first stage, a conditional random field (CRF) is used to calculate the probability that each gene in a user-supplied (meta)genome belongs to a secretion system; gene clusters that surpass a probability threshold are considered to be secretion system-encoding gene clusters. In the second stage, secretion system-encoding gene clusters (if present) are assigned to one or more of eight single-locus secretion system types (or are classified as “unknown” if a probability threshold is not surpassed for any type). **(B)** Receiver operating characteristic (ROC) and **(C)** precision-recall (PR) curves constructed via 10-fold cross validation (CV). Curves denote (i) the final CRF implemented in Sismis, which underwent feature selection (i.e., hyperparameters *c1* = 10, *W* = 20, *T* = 0.15; “Sismis CRF”, dashed purple line); (ii) the same CRF with no feature selection (i.e., *c1* = 10, *W* = 20, *T* = 1.0; “No feature sel.”, solid light blue line). Area-under-the-curve (AUC) values are reported in the legends. Violin plots comparing **(D)** real (wall-clock) time (in seconds) and **(E)** peak resident set size (RSS; in megabytes [MB]) per genome for (i) the final CRF implemented in Sismis, which underwent feature selection (i.e., hyperparameters *c1* = 10, *W* = 20, *T* = 0.15; “Sismis”, purple); (ii) the same CRF with no feature selection (i.e., *c1* = 10, *W* = 20, *T* = 1.0; “No feat. sel.”, light blue). Both CRFs were benchmarked on the complete set of species representative genomes from proGenomes v3 (*n* = 41, 777 genomes).

Using 10-fold cross validation (CV), we found that CRFs with hyperparameters *c1* = 10 and *W* = 20 (corresponding to the L1 regularization hyperparameter and sliding window size in number of genes, respectively) consistently outperformed other models (**Supplementary Figures S1-S2**). Notably, when we employed a feature selection approach (nested within CV folds to prevent overfitting), the CRF feature space could be reduced dramatically without noticeable drops in accuracy (i.e., CRFs with *c1* = 10, *W* = 20, and *T* = 0.15-1.0 produced similar results, where *T* corresponds to the proportion of features included in the model; **Figure 1BC**). Compared to a CRF with no feature selection employed (i.e., *T* = 1.0), the CRF with *T* = 0.15 was 3x faster and used less memory on average (**Figure 1DE**). We thus selected *c1* = 10, *W* = 20, and *T* = 0.15 as the final set of hyperparameters and implemented the corresponding model in Sismis. When applied to the full test set (*n* = 400 contigs) and conservative subset (*n* = 180 test set contigs with < 90 ANI relative to all training set contigs), the final Sismis CRF was able to detect secretion system gene clusters with high accuracy, achieving AUPR scores of 0.71 and 0.58, respectively (**Figure 2AB**).

**Figure 2.**
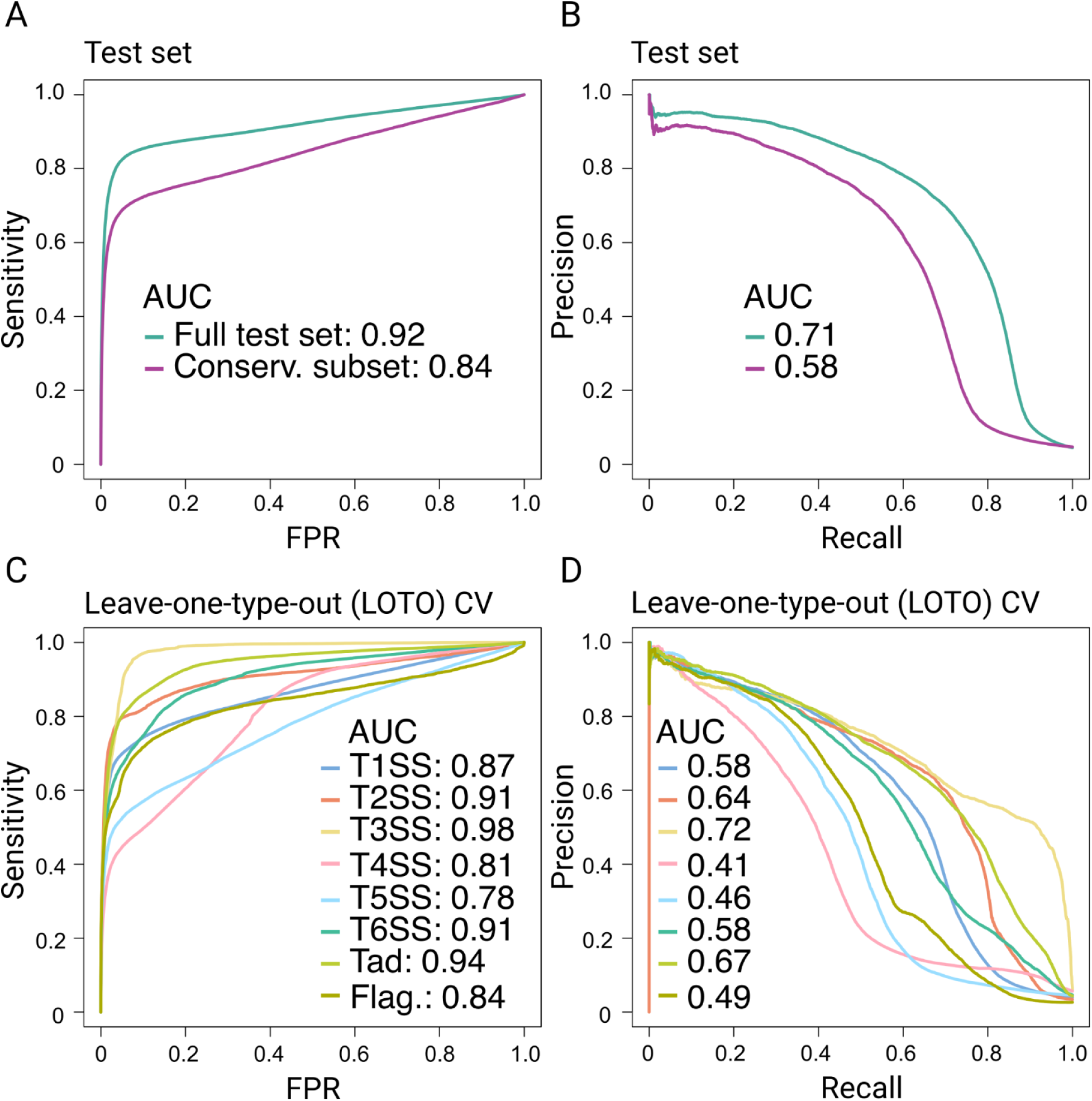
The Sismis CRF can be used for novel secretion system discovery tasks. **(A)** Receiver operating characteristic (ROC) and **(B)** precision-recall (PR) curves evaluating the performance of the final Sismis CRF (i.e., hyperparameters *c1* = 10, *W* = 20, *T* = 0.15) on an external test set. Curves denote the performance of Sismis on (i) the full test set (*n* = 400 contigs; “Full test set”, green curve) and (ii) a subset of test set contigs with < 90 average nucleotide identity (ANI) relative to all training set contigs (*n* = 180 contigs; “Conserv. Subset”, purple curve). Area-under-the-curve (AUC) values are reported in the legends. **(C)** ROC and **(D)** PR curves constructed via leave-one-type-out (LOTO) cross validation (CV). Line color denotes the secretion system type (per TXSSdb) used for validation, with AUC values reported in the legends. TXSS, Type X secretion system; Tad, tight adherence pili; Flag., Flagellum.

### Sismis identifies secretion system types on which it has not been trained

In addition to an external test set evaluation, we used leave-one-type-out (LOTO) CV to assess the performance of the final Sismis CRF on secretion systems of previously unseen types, simulating a novel secretion system discovery task. Within this framework, a CRF is trained on all TXSSdb secretion system types except one (the test set). Using LOTO CV, the final Sismis CRF achieved AUROC scores >0.80 for 7 of 8 single-locus secretion system types; for 4 types, AUROC scores >0.90 were achieved (Figure 2CD). The lowest AUROC score occurred for T5SS (AUROC = 0.78; Figure 2CD). While considerably smaller than other secretion system types in TXSSdb (i.e., they only span the cell outer membrane) (36), T5SS are diverse (e.g., they consist of at least five subclasses) (36, 37), and this may in part explain why T5SS are slightly more challenging to predict, having never encountered them before.

Comparatively, the highest AUROC and AUPR scores were both achieved for T3SS (0.98 and 0.72, respectively; Figure 2CD). One possible explanation for this could be that T3SS have similar genomic architecture to flagella (1, 38, 39), themselves one of the eight secretion system types in TXSSdb. Thus, T3SS may not pose a challenge to models that have encountered flagellar gene clusters before. Altogether, the LOTO CV approach used here provides insight into secretion system type discoverability and showcases the Sismis CRF’s strong overall performance at such a task.

### The Sismis CRF relies on annotated protein domains, as well as domains of unknown function (DUFs), for secretion system detection

A total of 678 protein domains were assigned positive (i.e., secretion system-associated) weights by the Sismis CRF, indicating that they are important for secretion system detection (Figure 3A). A GO enrichment analysis among positively weighted protein domains supported this, as GO terms associated with protein secretion and transport (e.g., GO:0009306, “protein secretion”; GO:0071806, “protein transmembrane transport”), secretion system activity (e.g., GO:0015628, “protein secretion by the type II secretion system”; GO:0007155, “cell adhesion”), and secretion system/flagellar structures (e.g., GO:0015627, “type II protein secretion system complex”; GO:0009288, “bacterial-type flagellum”) were among significantly enriched GO terms (topGO Fisher’s exact test raw *P*-value < 0.05; Figure 3B).

**Figure 3.**
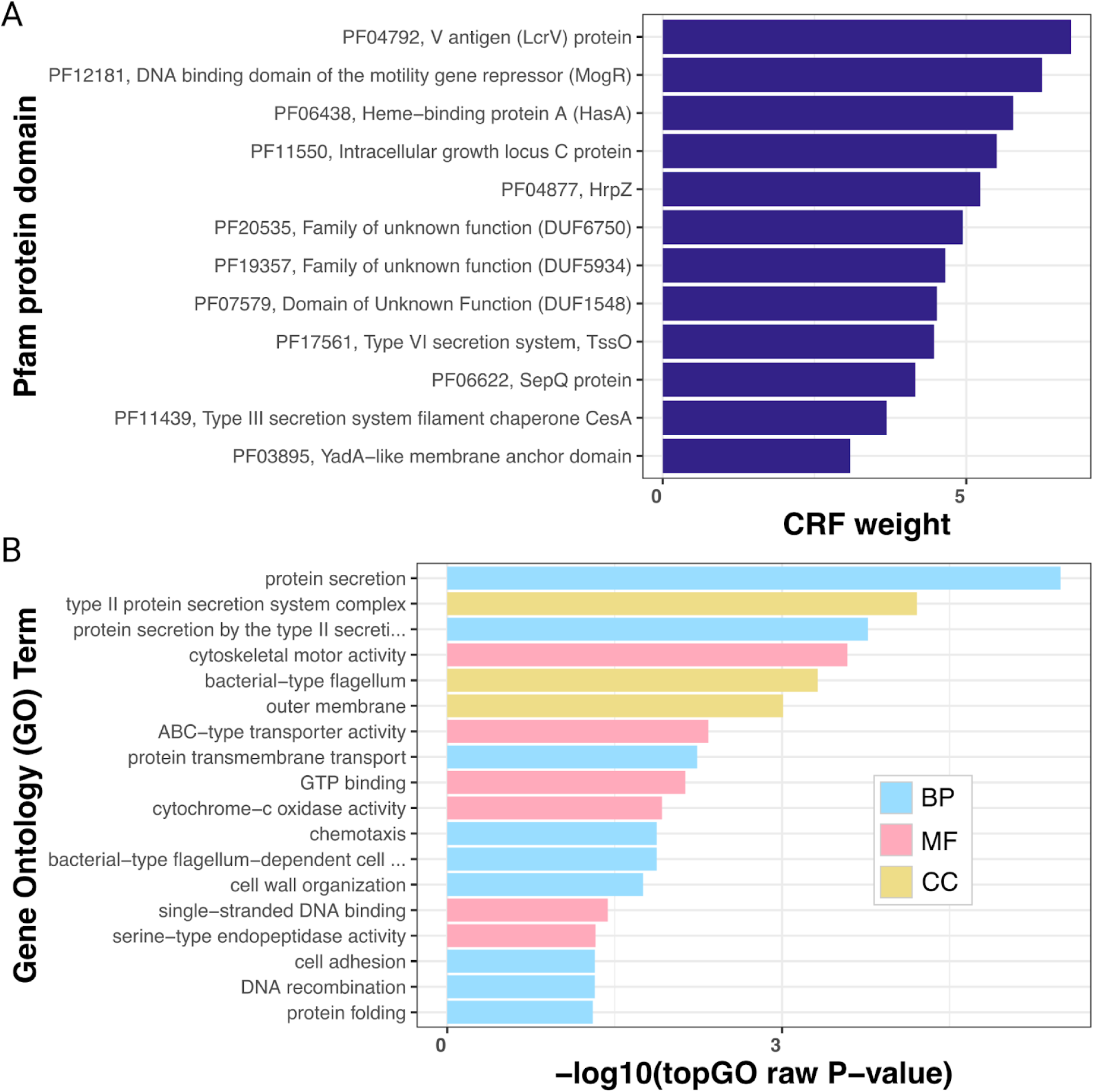
The Sismis CRF relies on annotated protein domains, as well as domains of unknown function (DUFs), for secretion system detection. **(A)** Protein domains (Y-axis) receiving the highest CRF weights (X-axis) in the final Sismis CRF (protein domains with weight ≥3.0 are shown for readability). **(B)** Gene Ontology (GO) terms (Y-axis) enriched among secretion system-associated (CRF weight > 0) protein domains in the Sismis CRF (topGO Fisher’s exact test raw *P* < 0.05; X-axis). BP, biological process; MF, molecular function; CC, cellular component.

At the individual protein domain devel, the five most highly weighted domains (CRF weight > 5.0) were: (i) V antigen LcrV (PF04792), a virulence factor secreted by the *Yersinia pestis* T3SS (40, 41), detected in multiple genera (e.g., *Yersinia, Vibrio, Pseudomonas, Aeromonas*); (ii) MogR (PF12181), a transcriptional repressor of flagellar genes in *Listeria monocytogenes* (42), detected primarily in the *Bacillus* and *Listeria* genera; (iii) HasA (PF06438), a hemophore secreted by *Serratia marcescens* (43), detected in multiple genera (e.g., *Pseudomonas, Yersinia, Serratia*); (iv) IglC (PF11550), secreted by the *Francicella* T6SS (44, 45), confined almost exclusively to the *Francicella* genus, but additionally detected in a genome of the recently discovered *Cysteiniphilum litorale* (*46, 47*); (v) HrpZ (PF04877), secreted and exported by the T3SS of plant pathogen *Pseudomonas syringae* (48), detected in multiple genera (e.g., *Pseudomonas*, *Pectobacterium, Dickeya, Erwinia*, per InterPro, https://www.ebi.ac.uk/interpro/search/text/, accessed 19 August 2025; Figure 3A).

Interestingly, 86 yet-uncharacterized domains of unknown function (DUFs) received positive weights in the Sismis CRF, including 3 DUFs with CRF weight > 4.0 (Figure 3A). Among these were PF20535, a DUF detected in multiple genera within Enterobacteriaceae (e.g., *Salmonella, Escherichia, Enterobacter, Klebsiella*) and elsewhere (e.g., *Pseudomonas, Serratia, Yersinia*). According to UniProt, this DUF is found in proteins annotated as e.g., “conjugal transfer protein” (e.g., TraR, TraO, TrbC), “IncI1-type conjugal transfer protein TraR”, “TrbC/VirB2 family protein”, “Integrating conjugative element membrane protein”, and “Type IV secretion protein IcmD” (https://www.uniprot.org/uniprotkb?query=PF20535, accessed 20 August 2025). This is notable, as some conjugative plasmids (e.g., IncI1) encode a T4SS (49–51). Further, IcmD has been shown to be required by *Coxiella burnetii* for T4SS secretion and productive infection of human macrophages (52, 53); similar type IVB secretion systems (T4BSS) are necessary for the pathogenesis of other organisms (e.g., *Legionella pneumophila*) (54, 55).

Comparatively, PF07579, also one of the highest-weighed DUFs, was exclusively present in *Chlamydia* spp. (per InterPro), in “uncharacterized” proteins, with no additional information available other than such proteins may be associated with the cell membrane (https://www.uniprot.org/uniprotkb?query=PF07579, https://www.uniprot.org/uniprotkb/Q5L578/entry, and https://www.uniprot.org/uniprotkb/Q5L577/entry, accessed 20 August 2025; Figure 3A). Together, these highly secretion system-associated DUFs may serve as potential leads for future experimental characterization and showcase the value of interpretable ML models.

### Sismis predictions from 700k (meta)genomes indicate that the vast majority of secretion systems are not represented in existing public databases

To showcase its scalability, we used Sismis to detect secretion systems across the entirety of the mOTUs v3 database (14), which we then clustered into gene cluster families (GCFs) alongside all TXSSdb secretion systems (*n* = 9, 756 TXSSdb secretion systems). Among the 694, 542 mOTUs (meta)genomes, Sismis detected a total of 747, 439 secretion systems, which, together with TXSSdb, could be partitioned into 15, 612 distinct GCFs (i.e., units that group similar secretion systems together; Figure 4).

**Figure 4.**
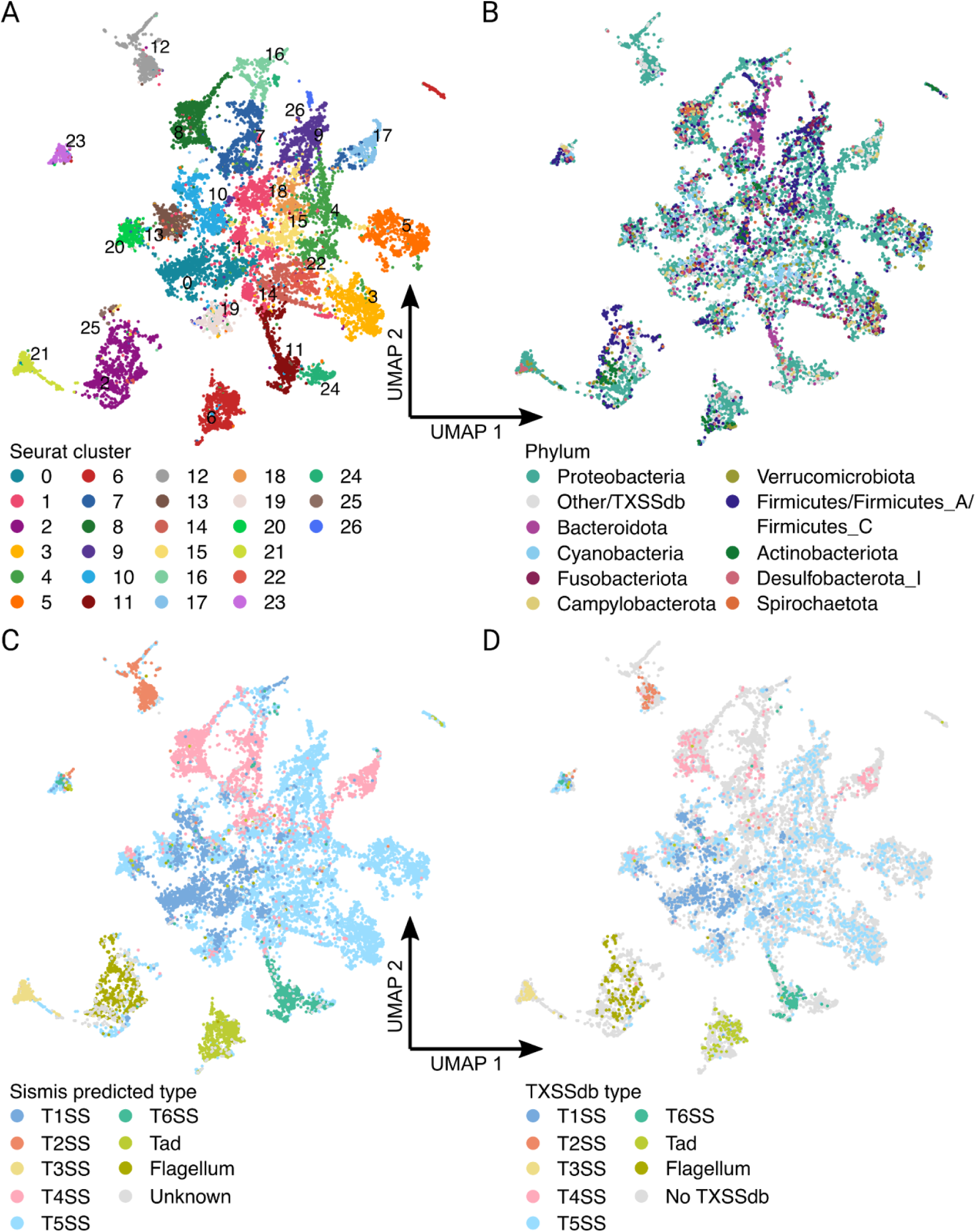
The Sismis Atlas contains predicted secretion systems from nearly 700k (meta)genomes. UMAP points denote gene cluster family (GCF) representatives, colored by **(A)** Seurat cluster (Seurat clustering threshold = 0.30); **(B)** phylum; **(C)** predicted secretion system type, obtained using the Sismis random forest classifier; **(D)** whether the GCF contains ≥1 TXSSdb secretion system (colors) or not (“No TXSSdb”, gray). For (B) and (C), points are colored by the majority class within the GCF. For (D), points with ≥1 TXSSdb secretion system are colored by the majority class within the GCF (only TXSSdb secretion systems are considered). GCFs with no TXSSdb secretion system (i.e., GCFs that contain Sismis predictions only) are represented by gray points (“No TXSSdb”).

A UMAP constructed using the protein domain composition of 15, 612 representative secretion systems (one per GCF) was partitioned into 27 Seurat clusters (Seurat clustering threshold = 0.30; Figure 4A). Secretion systems from Proteobacteria, the phylum in which over two-thirds of all Sismis secretion systems were detected (*n* = 505, 137 secretion systems, 67.6%), were distributed across 26 of 27 Seurat clusters (Figure 4AB); the remaining Seurat cluster (i.e., cluster 25) was nearly exclusive to Firmicutes/Firmicutes_A/Firmicutes_C and contained flagellar/predicted flagellar gene clusters (*n* = 54, 978 of 55, 065 Seurat cluster 25 secretion systems, 99.8%; Figure 4).

Within the secretion system UMAP, most secretion system types were confined to a single Seurat cluster (5 of 8 TXSSdb types, i.e., T2SS, T3SS, T6SS, Tad, Flagellum; Figure 4ACD). Specifically, considering “ground truth” types from TXSSdb only, 54 of 56 majority-T3SS GCFs (96.4%) were confined to a single Seurat cluster (Seurat cluster 21), with similar results observed for T2SS (52 of 56 GCFs in Seurat cluster 12; 92.9%), tight adherence (Tad) pili (107 of 117 GCFs in Seurat cluster 6; 91.5%), T6SS (105 of 126 GCFs in Seurat cluster 11; 83.3%), and flagella (131 of 168 GCFs in Seurat cluster 2; 78.0%, Figure 4AD). Majority-T1SS GCFs were largely (>70%) confined to two Seurat clusters (i.e., 334 and 102 of 610 GCFs in Seurat clusters 0 and 10; 54.8% and 16.7%, respectively), with majority-T4SS GCFs confined to three (126, 50, and 48 of 294 GCFs in Seurat clusters 8, 17, and 7; 42.9%, 17.0%, and 16.3%, respectively, Figure 4AD). Comparatively, T5SS-majority GCFs (*n* = 1, 481) were distributed across the secretion system UMAP (Figure 4AD).

Notably, of 15, 612 GCFs, the overwhelming majority (12, 704, 81.4%) did not contain any TXSSdb secretion systems (Figure 4D), indicating that Sismis detects secretion systems that are not present in TXSSdb. Of particular interest was Seurat cluster 6 (Figure 4AD); using an increased Seurat clustering threshold of 0.50, Seurat cluster 6 split into two distinct, stable subclusters in two-dimensional space: (i) a large subcluster, to which nearly all tight adherence (Tad) pili-majority GCFs were confined (referred to hereafter as the “major Tad subcluster”; *n* = 746 GCFs, including 105 of 117 TXSSdb Tad-majority GCFs, 89.7%); (ii) a small subcluster of 120 GCFs, 118 of which (98.3%) contained no TXSSdb secretion systems (referred to hereafter as the “minor unknown subcluster”; Figure 5A). A total of 61 protein domains, all of which were more common in the major Tad subcluster/less common in the minor unknown subcluster, differentiated the two subclusters (Seurat adjusted *P*-value < 0.05). Many of these subcluster markers had ATP-related functions (e.g., AAA ATPase domains), an observation that was further supported by GO enrichment results (topGO Kolmogorov-Smirnov test raw *P*-value < 0.05; Figure 5B). This is notable, as ATPase are key components of Tad and other type IV pili (56–58).

**Figure 5.**
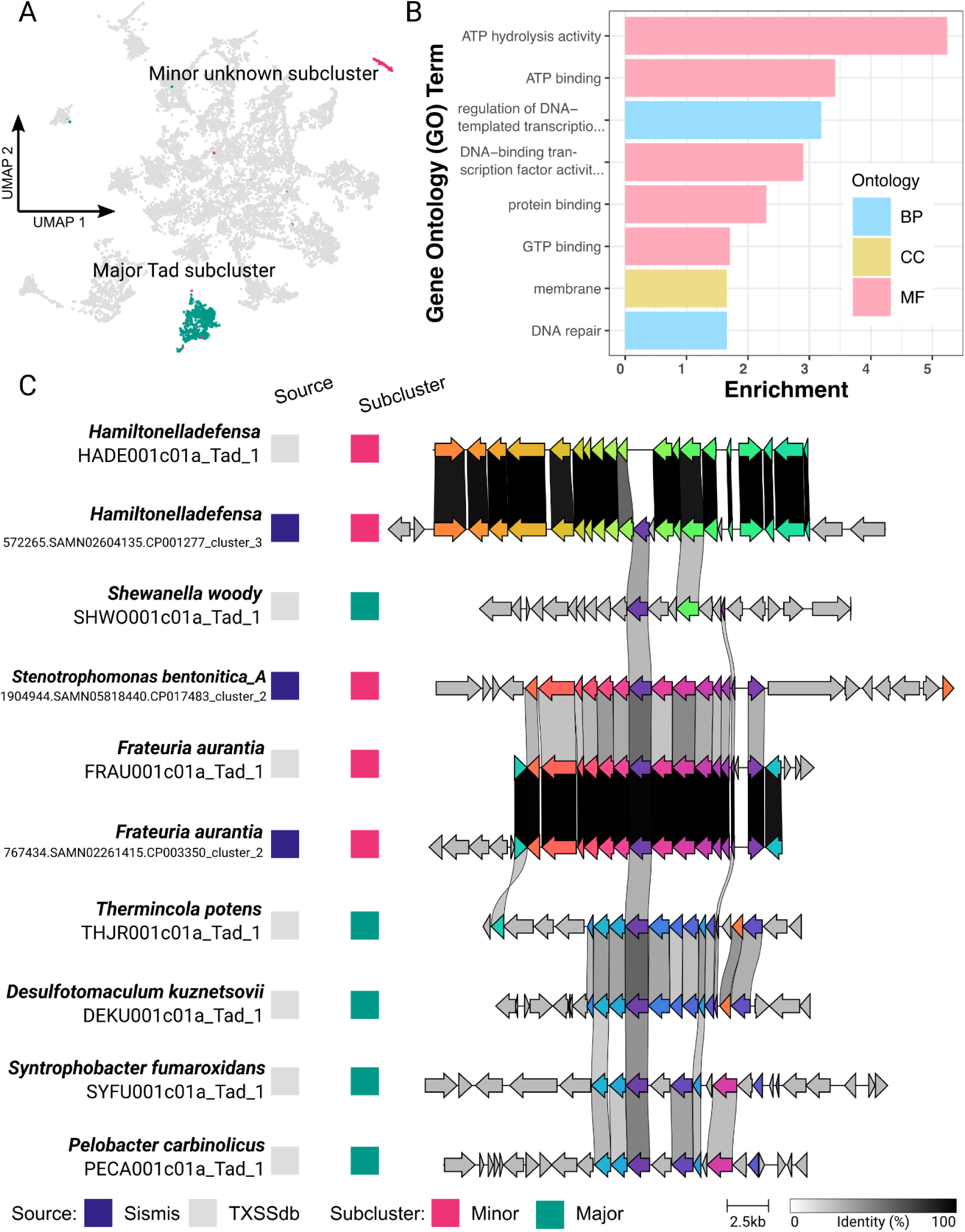
Sismis identifies a largely unexplored subcluster with tight adherence (Tad) pili-like characteristics. (A) Sismis Atlas UMAP, colored by subcluster (Seurat clustering threshold = 0.50): (i) a large subcluster with nearly all Tad pili-majority gene cluster families (GCFs; “Major Tad subcluster”, green, *n* = 746 GCFs); (ii) a small subcluster of 120 GCFs, 118 of which (98.3%) contained no TXSSdb secretion systems (“Minor unknown subcluster”, pink). Points denote GCF representatives. **(B)** Gene Ontology (GO) terms (Y-axis) enriched among protein domains that differentiate the major Tad subcluster from the minor unknown subcluster (topGO Kolmogorov-Smirnov test raw *P* < 0.05; X-axis). BP, biological process; MF, molecular function; CC, cellular component. **(C)** Comparison of secretion systems within the major Tad subcluster and minor unknown subcluster. The program clinker (https://github.com/gamcil/clinker) was used to compare secretion systems using default settings. Arrows denote genes within each secretion system gene cluster, with grayscale links denoting percent amino acid identity shared between corresponding genes. Colored boxes to the left of the secretion systems denote the source (Sismis or TXSSdb, “Source”) and subcluster (Minor or Major, “Subcluster”) of each secretion system. Text to the left of each secretion system denotes (i) the host species (top text) and (ii) unique identifier (bottom text) for each secretion system. For TXSSdb secretion systems, the species reported by TXSSdb is used. For Sismis secretion systems, Genome Taxonomy Database (GTDB) (59) species assignments available with mOTUs v3 were used.

Manual inspection of minor unknown subcluster secretion systems from complete genomes, plus their 50 Kbp flanking regions, alongside TXSSdb secretion systems from both subclusters confirmed minimal sequence similarity between the two subclusters (Figure 5C). The two sole TXSSdb secretion systems present in the minor unknown subcluster were each nearly identical to minor unknown subcluster secretion systems detected by Sismis (detected in *Hamiltonella defensa* and *Frateuria aurantia*; Figure 5C). Interestingly, Sismis identified a secretion system in a complete *Stenotrophomonas bentonitica_A* genome, which shared a low degree of amino acid identity with the *Frateuria aurantia* Tad secretion system (Figure 5C). This Tad-like secretion system is not included in TXSSdb, and to our knowledge, no such Tad-like secretion system has been described in *Stenotrophomonas bentonitica*. However, future experimental validation is needed to determine if this particular gene cluster, as well as other gene clusters in the minor unknown subcluster, actually exhibit secretion system-like functions, and if so, how members of this subcluster differ from canonical Tad systems (if at all).

To facilitate further exploration of secretion system data generated here, we integrated Sismis predictions from all 700k mOTUs v3 (meta)genomes into an interactive atlas (i.e., the Sismis Atlas; https://sismis.microbe.dev/). Written in Rust, the Sismis Atlas allows users to browse secretion system UMAPs (Figure 4), view/explore individual secretion system gene clusters, and download selected data (e.g., annotated secretion system gene clusters in GenBank format) and metadata (e.g., predicted type, host taxon). By making the results of our secretion system mining efforts publicly accessible, the Sismis Atlas can enable further large-scale secretion system exploration and discovery efforts.

## DISCUSSION

As sequential ML models, CRFs account for genomic context when making predictions (60). This, alongside the benefits that come with their inherent interpretability (61), make CRFs a natural fit for the annotation of structured prokaryotic gene clusters (e.g., secondary metabolite-associated biosynthetic gene clusters [BGCs] and prophage, as demonstrated previously) (10, 13). Here, we extend CRFs to the secretion system space and demonstrate that Sismis, our CRF-based tool, can detect secretion systems in prokaryotic (meta)genomes with high accuracy (full test set AUPR =0.71, AUROC = 0.92) and speed (using 1 CPU, <1 minute per genome on average).

While Sismis enables accurate secretion system annotation, it is important to point out its limitations. First, the CRF-based approach used here assumes that secretion systems are encoded by single gene clusters (termed single-locus systems per TXSSdb) (6). Sismis is not designed to detect secretion systems encoded by multiple loci spread throughout the genome (i.e., multi-locus systems; e.g., T9SS) (62). In the future, it may be possible to train accurate ML models that account for long contexts (e.g., long context language models) (63) to detect multi-locus secretion systems in prokaryotic (meta)genomes. However, such models would require a great deal of training data, likely far beyond what is available today. We thus recommend that users interested in multi-locus secretion system detection use designated tools/methods, which do not rely on the single gene cluster assumption (e.g., MacSyFinder, curated pHMMs) (8, 9).

Finally, it is important to note that there is no universally accepted definition for what constitutes a “secretion system” (3). Different definitions may include/exclude systems based on what is secreted (e.g., proteins, possibly alongside other macromolecules), where such molecules are secreted (e.g., across one or several bacterial membranes; out of the cell, possibly including molecules attached to the surface of the bacterium), and in which organisms (e.g., Gram-negative only; both Gram-negative and Gram-positive) (3). Secretion system type and subtype classification is similarly convoluted, with no clear definition for either (3). Thus, the performance of both rule- and ML-based secretion system detection and classification methods inherently depend on their definitions of secretion systems and their associated (sub)types. Here, we used secretion systems and (sub)types as defined by TXSSdb (6); however, some users may prefer a different secretion system classification framework.

Overall, Sismis and its companion atlas enable accurate secretion system annotation on an unprecedented scale. We anticipate that the methods/tools developed here will be particularly useful for e.g., large-scale secretion system mining initiatives, where computational efficiency is important; secretion system annotation in novel and/or under-studied taxa (i.e., those absent or under-represented in existing databases); and novel secretion system (sub)type discovery tasks, where rule-based approaches may struggle.

## DATA AVAILABILITY

Sismis is open-source and freely available via GitHub under the GNU General Public License v3 (https://github.com/lmc297/Sismis). Source code for the Sismis version used in the manuscript (v0.1.1) is available via Zenodo (https://doi.org/10.5281/zenodo.17019224). Source code for the Sismis Atlas is available via GitHub (https://github.com/henriksson-lab/sismis_atlas). Accession numbers of genomes/contigs used in the study, as well as all benchmarking and validation results generated in the study, are available via Zenodo (https://doi.org/10.5281/zenodo.17019224).

## SUPPLEMENTARY DATA

Supplementary Data are available via Zenodo (https://doi.org/10.5281/zenodo.17019224).

## AUTHOR CONTRIBUTIONS

Computational analyses were performed by LMC, FA, and JB. Software development was performed by ML, JH, and LMC. Dataset construction/curation was performed by ML and LMC. LMC conceived the study, which was funded by LMC and JH. LMC wrote the manuscript with contributions from all authors.

## ACKNOWLEDGMENTS

This research was conducted using the resources of High Performance Computing Center North (HPC2N; Umeå University, Umeå, Sweden). The Graphical Abstract and Figure 1 use an illustration from the NIAID NIH BioArt Source (https://bioart.niaid.nih.gov/bioart/123; “DNA”, Licensing: Public Domain).

## FUNDING

JB and LMC were supported by the SciLifeLab & Wallenberg Data Driven Life Science (DDLS) Program (grant KAW 2020.0239 to LMC), with additional funding provided by the Swedish Research Council (VR; grant 2023-05212 to LMC and grant 2024-06085 to JH/LMC). FA and JH were supported by a VR grant (grant 2024-03952 to JH), with additional funding provided by Cancerfonden (grant 23 3102 Pj to JH).

## CONFLICT OF INTEREST

The authors declare no competing interests.

## Notes

https://doi.org/10.5281/zenodo.17019224

### Competing Interest Statement

The authors have declared no competing interest.

